# Genome-scale CRISPR screening for modifiers of cellular LDL uptake

**DOI:** 10.1101/2020.07.01.182246

**Authors:** Brian T. Emmer, Emily J. Sherman, Paul J. Lascuna, Sarah E. Graham, Cristen J. Willer, David Ginsburg

## Abstract

Hypercholesterolemia is a causal and modifiable risk factor for atherosclerotic cardiovascular disease. A critical pathway regulating cholesterol homeostasis involves the receptor-mediated endocytosis of low-density lipoproteins into hepatocytes, mediated by the LDL receptor. We applied genome-scale CRISPR screening to query the genetic determinants of cellular LDL uptake in HuH7 cells cultured under either lipoprotein-rich or lipoprotein-starved conditions. Candidate LDL uptake regulators were validated through the synthesis and secondary screening of a customized library of gRNA at greater depth of coverage. This secondary screen yielded significantly improved performance relative to the primary genome-wide screen, with better discrimination of internal positive controls, no identification of negative controls, and improved concordance between screen hits at both the gene and gRNA level. We then applied our customized gRNA library to orthogonal screens that tested for the specificity of each candidate regulator for LDL versus transferrin endocytosis, the presence or absence of genetic epistasis with *LDLR* deletion, the impact of each perturbation on LDLR expression and trafficking, and the generalizability of LDL uptake modifiers across multiple cell types. These findings identified several previously unrecognized genes with putative roles in LDL uptake and suggest mechanisms for their functional interaction with LDLR.

## INTRODUCTION

Atherosclerotic cardiovascular disease is the leading cause of morbidity and mortality worldwide. A preponderance of evidence from epidemiology, human genetics, animal studies, and clinical trials have established that dysregulation of cholesterol homeostasis is a key factor in the pathogenesis of atherosclerosis[1]. Cholesterol is transported in the bloodstream in the form of lipoproteins, lipid-protein complexes that are typically characterized by their buoyancy during fractionation by ultracentrifugation. Cholesterol circulating in low-density lipoprotein (LDL) and other apolipoprotein B-containing lipoproteins exhibits a particularly strong correlation with atherosclerosis, and therapies that lower LDL cholesterol reduce the rate of cardiovascular disease. LDL cholesterol levels are tightly controlled through the complex interplay between intestinal absorption of dietary cholesterol, *de novo* cholesterol biosynthesis, efflux of cholesterol from peripheral tissues, and cellular uptake of lipoproteins[2].

A rich history of discovery in diverse fields including genetics, cell biology, and biochemistry has elucidated many of the molecular determinants of LDL regulation[3–5]. LDL is cleared from circulation by the LDL receptor (LDLR). The extracellular domain of LDLR directly binds to the apolipoprotein B component of LDL particles, triggering the receptor-mediated endocytosis of the LDLR-LDL complex into clathrin-coated vesicles. Internalized complexes of LDL and LDLR traffic through the endolysosomal pathway until luminal acidification triggers their dissociation, with cholesterol being extracted from LDL while LDLR either recycles back to the cell surface or, if bound to its negative regulator PCSK9, traffics to lysosomes for degradation. The importance of LDL uptake to human cholesterol regulation and cardiovascular disease is highlighted by the monogenic causes of familial hypercholesterolemia that affect this pathway[6], including mutations in the genes encoding for LDLR itself, its ligand apolipoprotein B, its negative regulator PCSK9, or its endocytic adapter LDLRAP1. An additional level of regulation of the genes involved in cholesterol uptake and synthesis is provided by SREBP signaling, in which low cellular sterol levels lead to the SCAP-mediated trafficking of SREBP proteins from the ER to the Golgi, where they are cleaved by resident proteases (encoded by *MBTPS1* and *MBTPS2*) to release and activate their transcription factor domains[7, 8]. Human genome-wide association studies (GWAS) have also identified >250 loci associated with blood lipid levels[9–11]. Despite these many successes, our molecular understanding of LDL regulation remains incomplete. For the majority of GWAS associations, the causal link to a specific gene and the mechanism for the genotype-phenotype correlation remains unknown. Moreover, only an estimated 20-30% of the heritability of lipid traits is currently explained[12]. It is therefore likely that additional, as yet unrecognized genetic interactions contribute to cholesterol regulation in humans.

Recent advances in genome editing and massively parallel DNA sequencing have enabled high-throughput functional interrogation of the mammalian genome[13]. We previously performed a genome-wide CRISPR screen for the molecular determinants of PCSK9 secretion, leading to our identification of SURF4 as a cargo receptor that recruits PCSK9 into the secretory pathway[14]. We now report adaptation of this approach to screen for modifiers of cellular LDL uptake. Through a primary genome-wide CRISPR screen, followed by the synthesis and re-screening of a focused secondary gRNA library with greater depth of coverage, we validated 118 positive regulators and 45 negative regulators of HuH7 cell LDL uptake. Orthogonal screening, in which this customized guide RNA (gRNA) library was applied to other phenotypic selections, enabled further characterization of putative hits for their specificity in influencing the endocytosis of LDL, the nature of their interaction with LDLR, and their generalizability across cell types.

## RESULTS

### Primary genome-scale CRISPR screen for modifiers of HuH7 cell LDL uptake

HuH7 cells, originally derived from a well-differentiated human hepatocellular carcinoma, are widely used as a model of hepatocyte LDL uptake[15]. We first investigated the time- and dose-dependence of LDL uptake by HuH7 cells. LDL uptake by HuH7 cells was readily detectable above cellular autofluorescence by flow cytometry following a 1 hour incubation with 4 μg/mL of fluorescently-conjugated LDL in serum-free media (Supplemental Figure 1A). These conditions were in the linear range of detection with respect to both LDL dose and duration of incubation (Supplemental Figure 1B-C). Acquisition of fluorescent signal was mediated by LDLR, as CRISPR-mediated targeting of *LDLR* resulted in a ~75% reduction in LDL uptake under these conditions (Supplemental Figure 1D). Pre-incubation of HuH7 cells with lipoprotein-depleted media resulted in an *LDLR*-dependent ~67% increase in LDL uptake, consistent with upregulation of *LDLR* expression via SREBP signaling (Supplemental Figure 1D). These results suggest that this model system recapitulates the LDLR-dependence and SREBP-responsiveness of cellular LDL uptake and is a suitable platform for genome-wide screening.

To comprehensively identify genetic modifiers of HuH7 cell LDL uptake, we transduced ~25 million cells with the pooled GeCKOv2 lentiviral library delivering Cas9 and 123,411 gRNAs, including 6 gRNA for nearly all known protein-coding genes in the genome[16] (Figure 1A). Following 13 days of expansion in culture, to facilitate target site mutagenesis and turnover of wild-type protein, cells were split and cultured for an additional 1 day, either under continued lipoprotein-rich or changed to lipoprotein-depleted growth conditions. Mutant cells were then incubated with fluorescently-conjugated LDL and sorted by flow cytometry into bins of LDL^high^ (top 7.5%) and LDL^low^ (bottom 7.5%) cells. Massively parallel sequencing of amplified gRNA sequences from each bin was performed and the relative enrichment of each gRNA in either pool was assessed. A total of 3 independent biologic replicates were performed for each screen.

**Figure 1.**
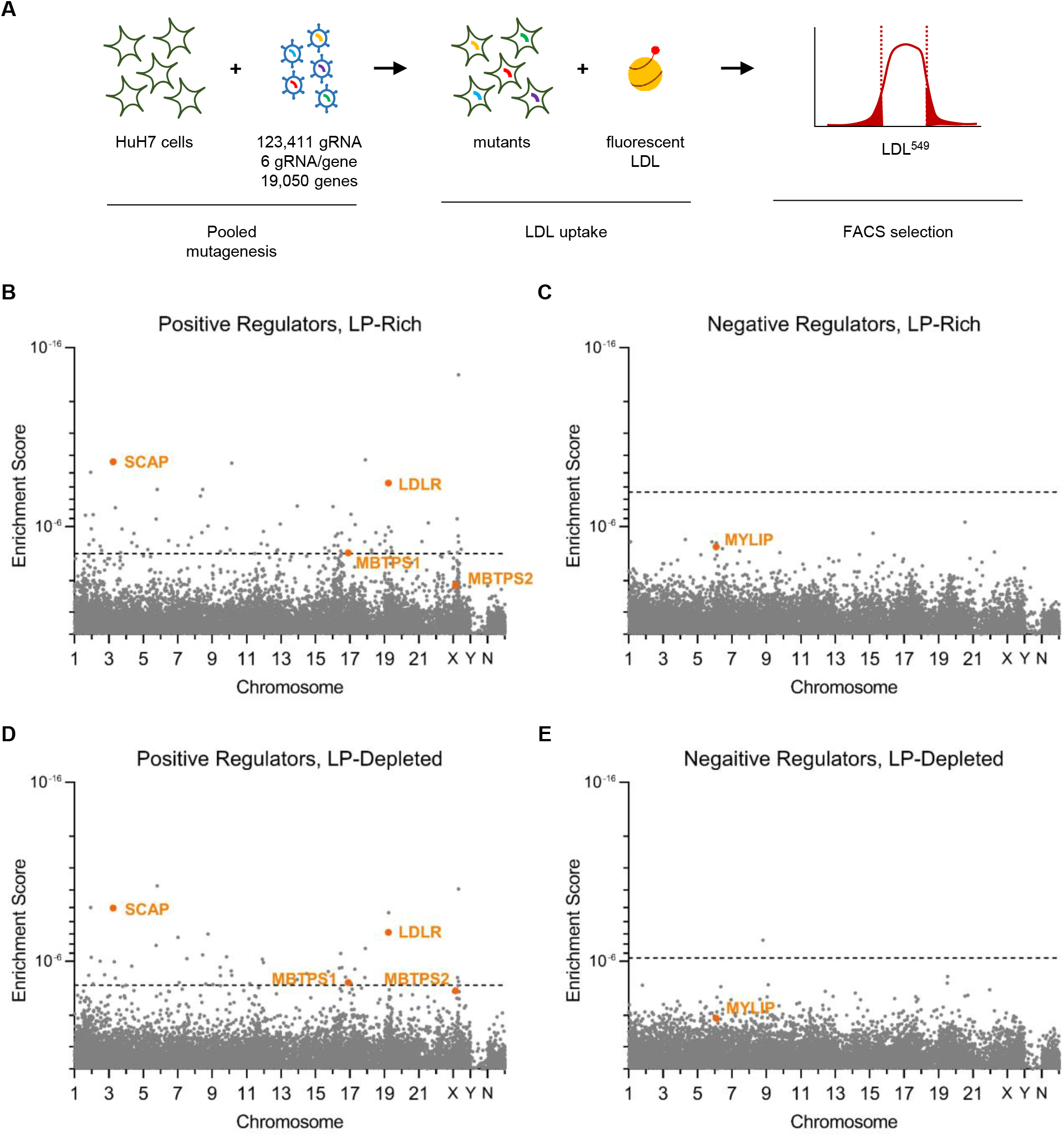
Primary genome-wide CRISPR screens for HuH7 LDL uptake modifiers. (A) Schematic of screening strategy. (B-E) MAGeCK gene level enrichment scores for genes who perturbation causes reduced LDL uptake (B, D) or increased LDL uptake (C, E) under lipoprotein-rich (B-C) or lipoprotein-depleted (D-E) culture conditions.

Gene-level analysis identified 95 candidate genes with a false discovery rate (FDR) <5% whose targeting was associated with reduced LDL uptake under either lipoprotein-rich or lipoprotein-deficient conditions (Figure 1B-C, Supplemental Table 1). Among these candidates were known regulators of LDL uptake including *LDLR*, *SCAP*, and *MBTPS1*, though *MBTPS2* was not identified among the screen hits (ranking 343 and 58 under lipoprotein-rich and lipoprotein-depleted conditions, with FDR>5% for both). A high degree of concordance was observed for identified positive regulators between the screens conducted under lipoprotein-rich or lipoprotein-depleted conditions, with 27/95 hits identified under both conditions (Supplemental Figure 2). Genes positively identified under lipoprotein-rich conditions were 246-fold more likely than negative genes to also be identified under lipoprotein-depleted conditions; genes positively identified under lipoprotein-depleted conditions were 384-fold more likely than negative genes to also be identified lipoprotein-rich conditions.

Only 1 gene, *SQLE*, was identified whose targeting was associated with enhanced LDL uptake (Figure 1D-E, Supplemental Table 1). The positive control *MYLIP*, encoding the LDLR negative regulator IDOL[17], was ranked 7 and 138 among negative regulators under lipoprotein-rich and lipoprotein-depleted conditions but did not meet the FDR<5% threshold for genome-wide significance. The identification of several positive controls and the concordance of our hits across screening conditions suggested that our primary screen was successful, though limited by background noise at genome scale.

### Secondary screen validation of HuH7 LDL uptake modifiers

To validate and refine our primary screen hits, we next developed a focused secondary screen of candidate genes with greater depth of coverage. We applied our validation testing to an extended list of potential LDL uptake regulators (positive regulators FDR<50%, negative regulators FDR<75%), reasoning that false negatives might lie further down our candidate list due to a variety of factors including inadequate gRNA efficiency or depth of coverage in the primary screen. We designed and synthesized a custom CRISPR library containing 12,207 gRNAs, including 15 gRNA per gene for 554 potential positive regulators and 170 potential negative regulators, along with 1000 control non-targeting sequences. Massively parallel sequencing of the plasmid pool of this library confirmed the presence of 99.97% of gRNA sequences inserted into the CRISPR plasmid backbone, (Supplemental Figure 3A) with minimal library skewing (Supplemental Figure 3B). We generated lentiviral pools from this plasmid mix and performed a secondary screen for HuH7 cell LDL uptake using conditions that were identical to our primary screen, aside from greater depth of coverage at all stages owing to the smaller library size (Supplemental Figure 4). Using an FDR cutoff of 5% in our secondary screen, we identified 118 positive regulators of HuH7 LDL uptake (Figure 2A-B, Supplemental Table 2), with 66 of these exhibiting significant effects under both lipoprotein-rich and lipoprotein-depleted conditions (2C). We also identified 45 negative regulators, with 18 of these exhibiting significant effects under both lipoprotein-rich and lipoprotein-depleted conditions (Figure 2D). The validation rate of candidates in the secondary screen was strongly correlated to the strength of signal in the primary screen (Figure 2E). As in the primary screen, genes identified under either lipoprotein-rich or lipoprotein-depleted conditions were much more likely to be identified under the other condition (Supplemental Figure 5A-B), with a high degree of correlation for the relative effect size under either condition (Figure 2F). This concordance between screen conditions also extended to the individual gRNA level, as the relative ranking (Figure 2G) and magnitude of enrichment (Supplemental Figure 5C) for individual gRNAs under lipoprotein-rich conditions was correlated with their corresponding value under lipoprotein-depleted conditions.

**Figure 2.**
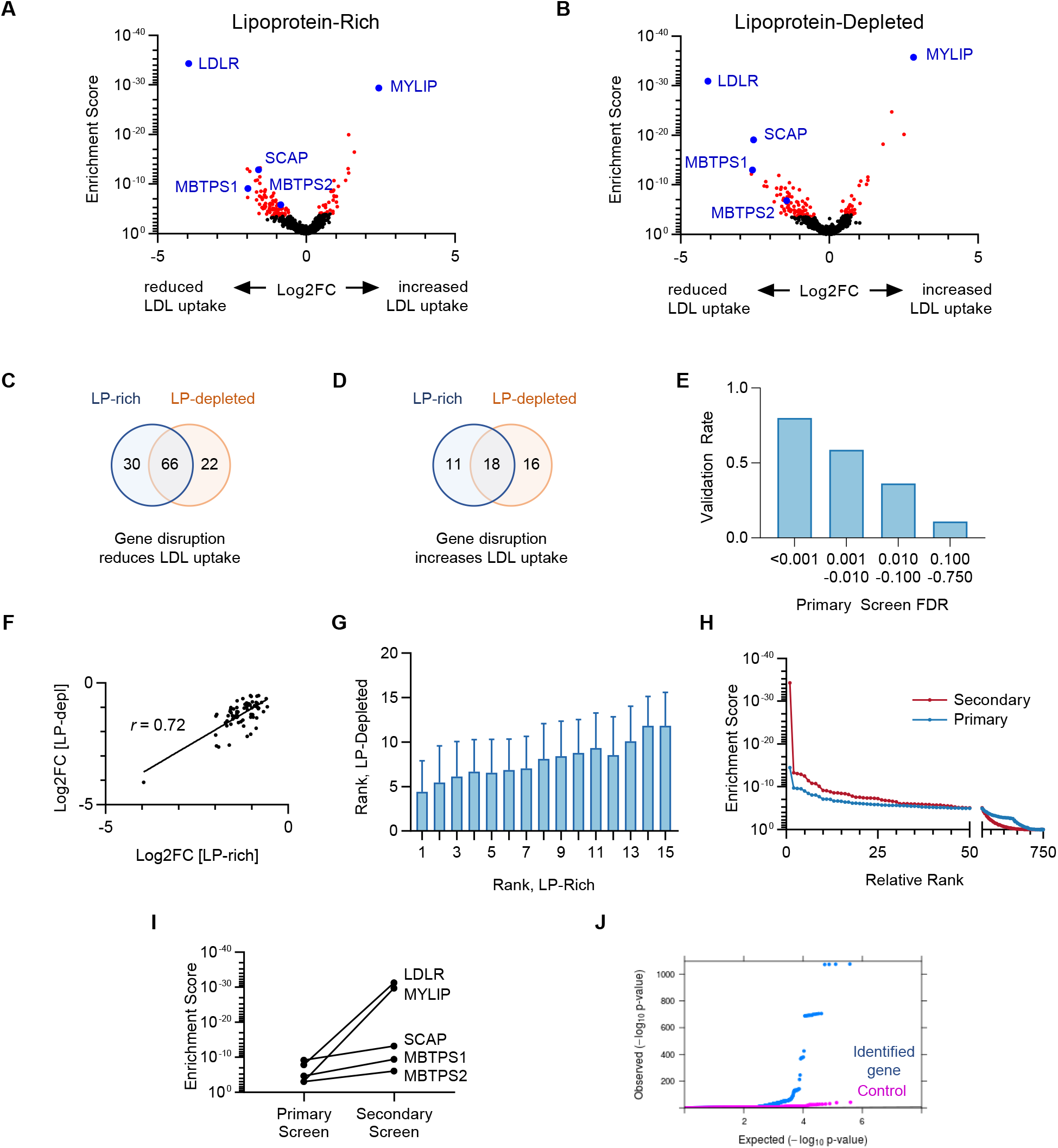
Targeted secondary CRISPR screens for modifiers of LDL uptake by HuH7 cells. (A-B) Volcano plots displaying MAGeCK gene level enrichment scores and associated gRNA log2 fold changes for each gene tested in the secondary gRNA library, under lipoprotein-rich (A) or lipoprotein-depleted (B) cultured conditions, with genes identified with FDR<5% displayed in red and positive controls in blue. (C-D) Venn diagrams of genes identified whose targeting was associated with reduced (C) or enhanced (D) cellular LDL uptake under lipoprotein-rich and/or lipoprotein-depleted culture conditions. (E) Genes identified in the primary screen for LDL uptake were stratified by FDR tier and compared for their validation rate (FDR<5%) in the secondary screen for LDL uptake. (F) Correlation of effect size for genes identified as positive regulators of LDL uptake under both lipoprotein-rich and lipoprotein-depleted culture conditions. (G) Cumulative distribution function of MAGeCK enrichment scores for genes tested in both the primary and secondary CRISPR screens for LDL uptake. (H) Comparison of MAGeCK gene level enrichment scores for positive control genes in the primary versus secondary CRISPR screens for LDL uptake. (J) QQ plot of LDL GWAS results in UK Biobank within identified LDL uptake regulator genes compared to matched control genes.

In accordance with its greater depth (both in terms of gRNA per gene tested, and cells per gRNA tested), the secondary screen exhibited more robust performance than the primary screen. More genes were identified with FDR<5%, suggesting an increased power of detection. Screen hits exhibited a clearer discrimination from genes with no signal (Figure 2H). Positive control genes *LDLR*, *SCAP*, and *MBTPS1* were again positively identified in the secondary screen, while *MBTPS2* and *MYLIP* (negative in the primary screen) were readily detected as positive hits in the secondary screen. Each of these internal control genes was identified with more significant enrichment (Figure 2I) and rose in the relative rankings, with *LDLR* and *MYLIP* becoming the top hits for reduced and enhanced LDL uptake, respectively, both under lipoprotein-rich and lipoprotein-depleted culture conditions.

### HuH7 LDL uptake regulators are enriched for LDL GWAS associations

Ontology analysis of our validated HuH7 LDL uptake regulators revealed significant enrichment for several annotations including genes involved in regulation of gene expression, cholesterol metabolism, Golgi to plasma membrane transport, protein N-linked glycosylation, and ubiquitin-mediated protein degradation (Supplemental Figure 6, Supplemental Table 3). Comparison to current human GWAS data from UK Biobank showed a significant enrichment for genes in proximity to genetic variants associated with LDL cholesterol relative to matched control genes. A total of 163 genes were identified to be either positive or negative regulators of HuH7 LDL uptake. Of these, 12% (20/163) had a genome-wide significant GWAS result (p-value < 5×10^−8^) within the gene while 33% (54/163) had a significant result within 500 kb. P-values for association with LDL cholesterol within the 163 identified genes were also more significant, on average, than those within length-matched control genes (two-sided p-value < 2.2×10^−16^, Figure 2J, Supplemental Table 4). The majority of our screen hits had not previously been implicated in cholesterol regulation.

### Most LDL uptake regulators do not cause a similar influence on transferrin uptake

To assess for nonspecific effects on global endocytosis, we next applied our customized gRNA library to assess HuH7 uptake of fluorescently-conjugated transferrin. *TFRC* was included among the secondary library gRNA target genes as a positive control. As expected, *TFRC* was the top hit whose disruption was associated with reduced transferrin uptake (Figure 3A, Supplemental Table 5). Among the 736 genes tested with our secondary library, 24 were found to positively regulate and 19 to negatively regulate transferrin uptake (FDR<0.05). Little concordance was observed between regulators of LDL and transferrin uptake (Figure 3B-D). Surprisingly, disruption of several genes resulted in decreased LDL uptake but enhanced transferrin uptake. Thus, the majority of hits from our secondary screen do not appear to result from global disruption or stimulation of receptor-mediated endocytosis.

**Figure 3.**
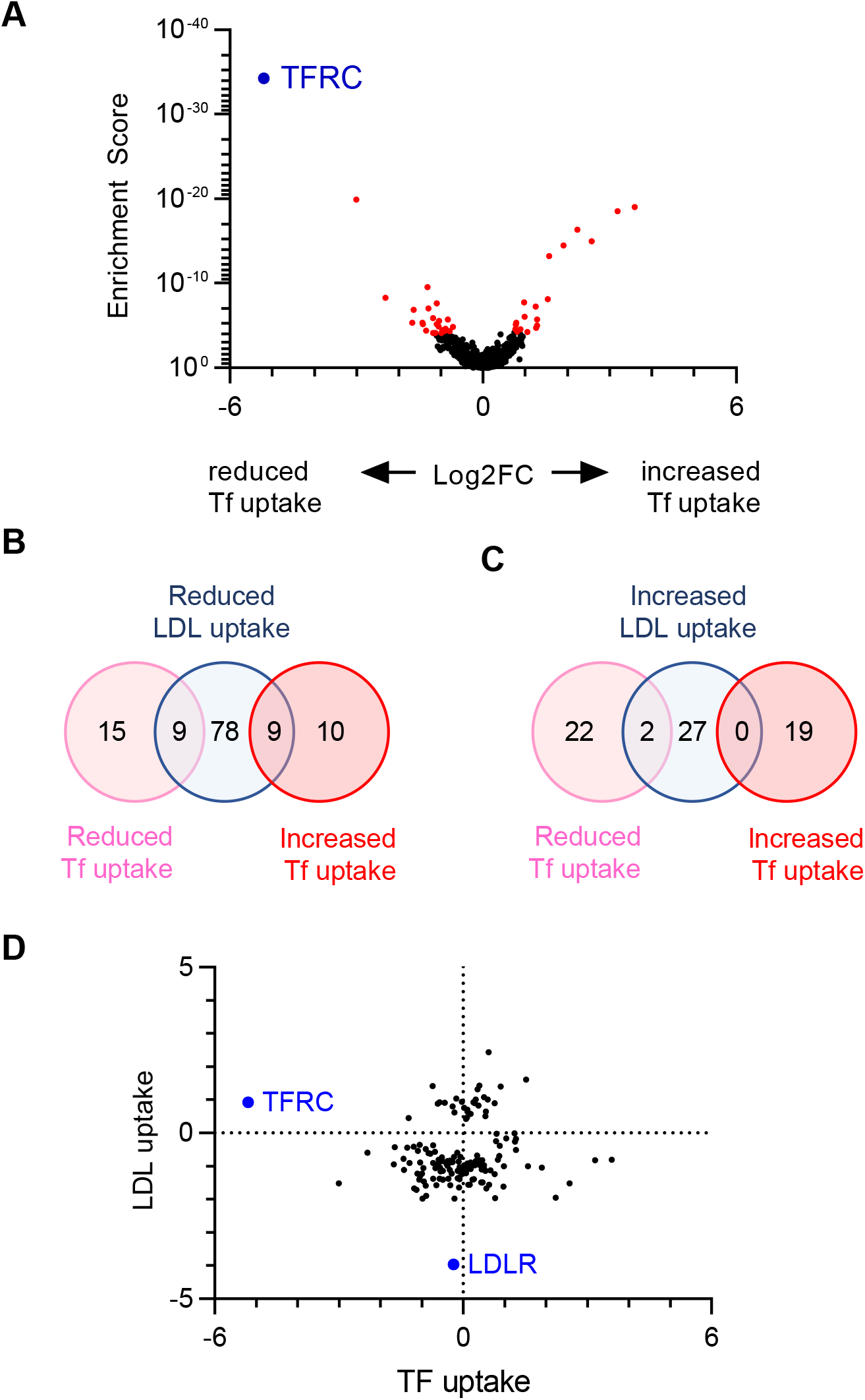
Orthogonal CRISPR screen for modifiers of transferrin uptake by HuH7 cells. (A) Volcano plot displaying transferrin uptake MAGeCK gene level enrichment scores and log2 fold change for each gene tested in the customized gRNA library, with genes identified with FDR<5% displayed in red and *TFRC* in blue. (B-C) Venn diagrams of genes identified whose targeting was associated with reduced (B) or increased (C) cellular transferrin uptake, in comparison to the effect of targeting each gene on HuH7 LDL uptake. (D) Relative effect sizes with log2 fold change for targeting of each gene on transferrin and LDL uptake.

### A subset of regulators influence LDL uptake independently of LDLR

Since the majority of fluorescent LDL acquisition under our screening conditions was *LDLR*-dependent (Supplemental Figure 1D), we hypothesized that most of our screen hits would influence LDL uptake via interaction with LDLR, and therefore would exhibit no effect on LDL uptake when tested on a *LDLR*-deleted genetic background. To test this hypothesis, we generated an HuH7 clone harboring a homozygous frameshift mutation in *LDLR* (Figure 4A), with no detectable LDLR protein by immunoblotting (Figure 4B) and a ~85% reduction in LDL uptake relative to parental wild-type cells (Figure 4C). We then screened this *LDLR*-deleted cell line with our secondary CRISPR library under lipoprotein-rich and lipoprotein-depleted culture conditions (Figure 4D-E, Supplemental Table 6). Surprisingly, we found that many modifiers of LDL uptake identified in wild-type cells were also identified in *LDLR*-deleted cells, with 45/118 positive and 17/45 negative regulators exhibiting similar effects on LDL uptake in *LDLR*-deleted cells (Figure 4F-I). This association is unlikely to be due to an influence on residual *LDLR* expression, as *LDLR*-targeting gRNAs were not enriched in LDL^low^ cells. Instead, these findings suggest that a significant subset of the LDL uptake modifiers identified here may influence LDLR-independent, or both LDLR-dependent and independent, LDL uptake.

**Figure 4.**
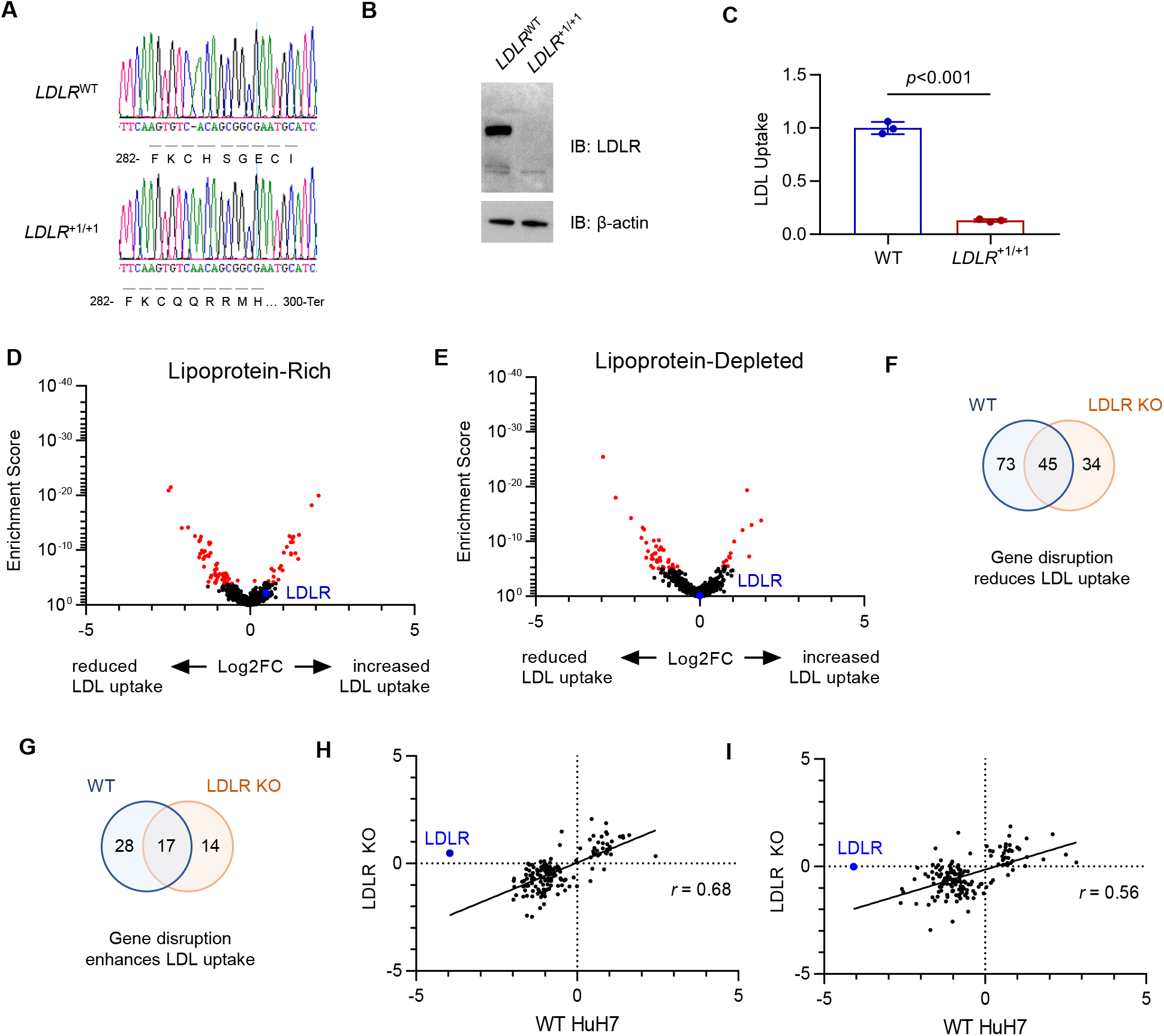
Orthogonal CRISPR screen for modifiers of LDL uptake by *LDLR*-deleted HuH7 cells. (A) Genotyping at the genomic DNA target site, (B) immunoblotting, and (C) quantification of LDL uptake by flow cytometry for a single cell HuH7 clone targeted by CRISPR at the *LDLR* locus. (D-E) Volcano plots displaying MAGeCK gene level enrichment scores and log2 fold change for each gene tested in the secondary gRNA library, under lipoprotein-rich (D) or lipoprotein-depleted (E) cultured conditions, with genes identified with FDR<5% displayed in red. (F-G) Venn diagrams demonstrating the overlap in genes identified from HuH7 WT and *LDLR* KO cells for genes whose disruption was associated with reduced (F) or enhanced (G) LDL uptake. (H-I) Comparison of effect size on LDL uptake in WT and *LDLR* KO cells under lipoprotein-rich (H) or lipoprotein-depleted (I) conditions for each gene showing a significant effect in either background.

### A subset of LDL uptake regulators modulate steady-state LDLR expression and trafficking to the cell surface

To determine how each of our screen hits influences LDLR activity, we mutagenized HuH7 wild-type cells with our customized gRNA library and selected mutants by the amount of LDLR staining either at the cell surface (Figure 5A-B, Supplemental Table 7) or in semi-permeabilized cells (Figure 5C-D, Supplemental Table 7). As expected, the top hit associated with both decreased surface and decreased total LDLR was *LDLR* itself, and the top hit for increased surface and increased total LDLR was *MYLIP*. We identified 26 and 20 genes whose targeting either reduced or enhanced LDLR surface staining, respectively (FDR<0.05, Figure 5A). Screening for total LDLR similarly revealed 46 and 43 genes whose targeting either reduced or enhanced total cellular LDLR staining (FDR<0.05, Figure 5C). Most targeted genes exhibiting decreased LDLR staining (surface or total cell-associated) had also exhibited decreased fluorescent LDL uptake (Figure 5E). In contrast, gene targeted cells with increased surface or total LDLR exhibited heterogeneous effects on LDL uptake, with roughly equal numbers exhibiting either reduced or increased LDL uptake (Figure 5E). Targeted genes demonstrated a high degree of correlation between surface and total LDLR staining (Figure 5F), with no genes exhibiting significant effects on surface and total LDLR staining in opposite directions.

**Figure 5.**
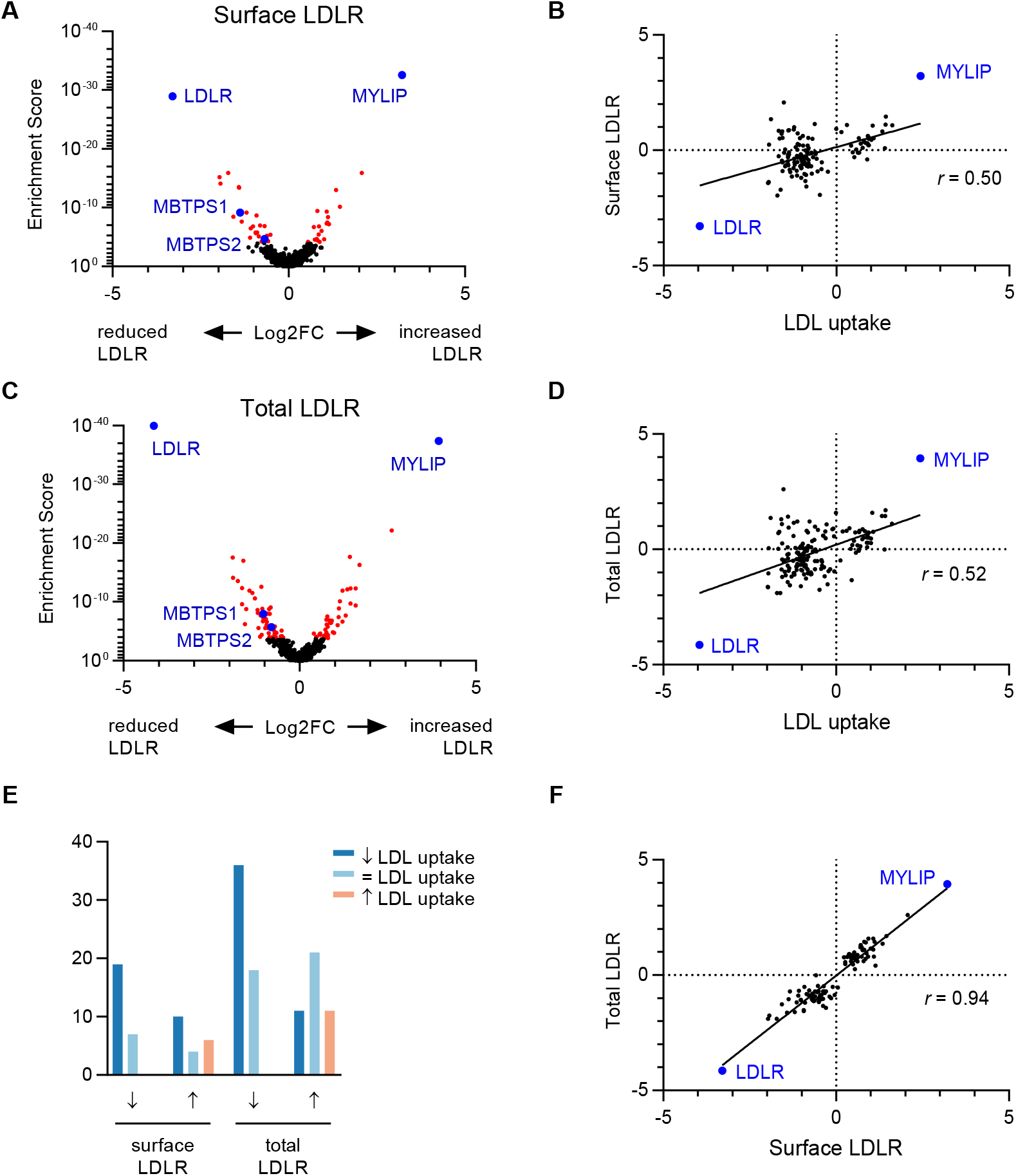
Orthogonal CRISPR screen for modifiers of LDLR abundance in HuH7 cells. (A) Volcano plot displaying surface LDLR abundance MAGeCK gene level enrichment score and log2 fold change for each gene tested in the customized gRNA library, with genes identified with FDR<5% displayed in red. (B) Comparison of effect size for LDL uptake and surface LDLR abundance for each gene showing a significant effect for either. (C) Volcano plot and (D) comparison of effect size for LDL uptake and total cellular LDLR abundance. (E) Comparison of corresponding effect on LDL uptake for each gene exhibiting an influence on surface or total LDLR abundance. (F) Comparison of effect size for each gene exhibiting an influence on either surface or total LDLR abundance.

### Cell-type specificity of LDL uptake modifiers

Comparison of our data to a previous siRNA screen for endothelial cell LDL uptake{Kraehling, 2016 #856} revealed limited overlap, with only 1 gene identified in both studies (Supplemental Table 8). To examine whether the LDL uptake modifiers identified here might be unique to HuH7 cells, we also applied our customized library to a screen of LDL uptake in HepG2 cells. As in HuH7 cells, LDL uptake in HepG2 cells was dependent on *LDLR* and modulated by targeting of known regulators including *SCAP*, *MBTPS1*, *MBTPS2*, and *MYLIP* (Figure 6A, Figure 6D, Supplemental Table 9). Under lipoprotein-rich conditions, we identified only 10 and 2 genes whose targeting was associated with reduced or increased LDL uptake in HepG2 cells, respectively (Figure 6A), with 6/10 positive regulators (Figure 6B) and 1/2 negative regulators (Figure 6C) exhibiting similar effects in HuH7 cells. A much higher number of LDL uptake modifiers were identified under lipoprotein-depleted conditions, with disruption of 53 and 5 genes associated with reduced or increased LDL uptake, respectively (Figure 6D volcano). Among these latter genes, 25/53 positive regulators (Figure 6E) and 2/5 negative regulators (Figure 6F) exhibited similar effects in HuH7 cells. The likelihood of a gene showing a functional influence on LDL uptake by HepG2 cells was predicted by the strength of its association with LDL uptake by HuH7 cells (Figure 6G). Significant positive correlation was observed for the degree of enrichment for a given LDL uptake modifier in either cell line (Figure 6H). No genes were identified that associated with significant effects on LDL uptake in opposite directions in HuH7 and HepG2 cells.

**Figure 6.**
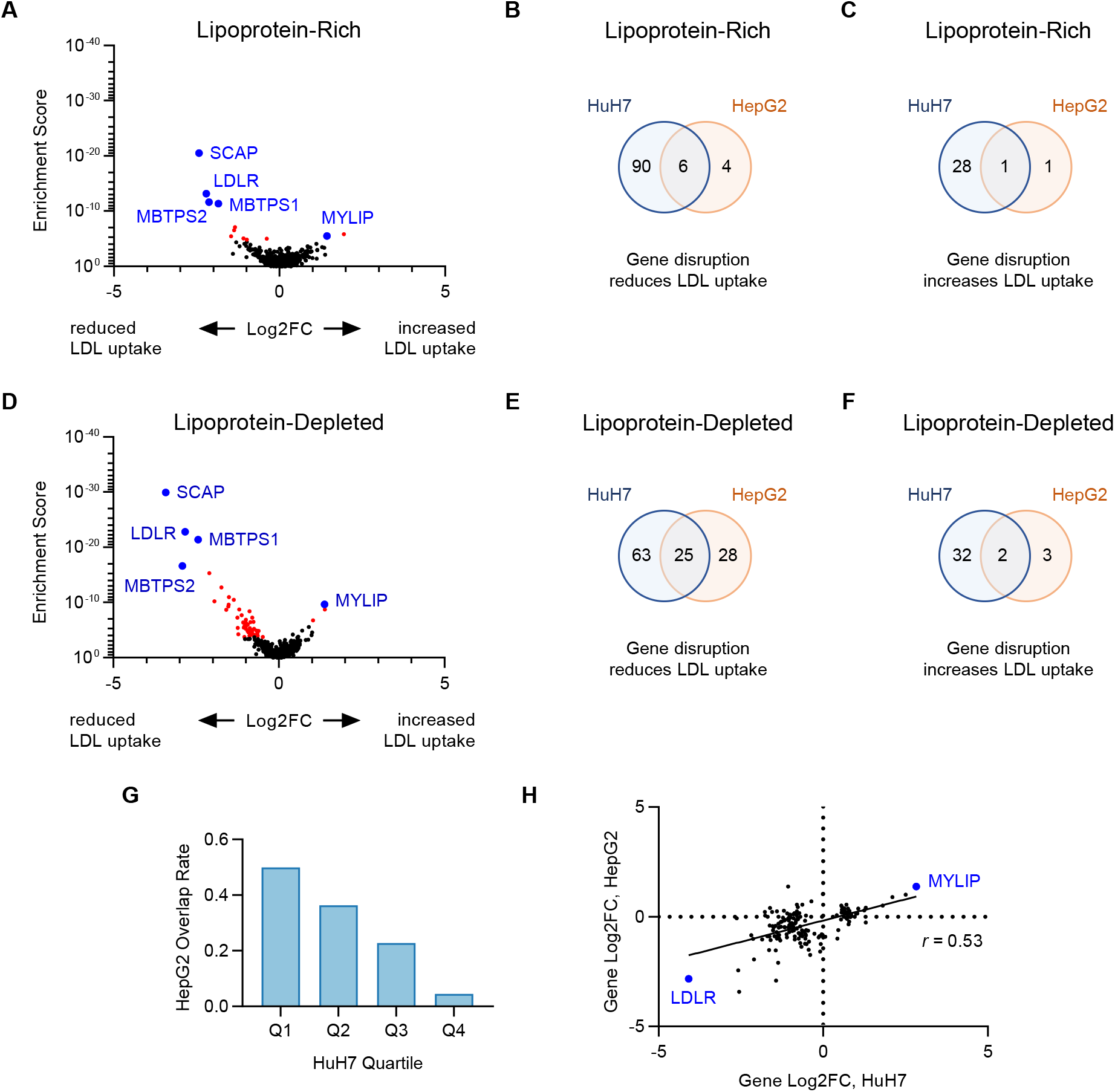
Orthogonal CRISPR screen for modifiers of LDL uptake by HepG2 cells. (A-F) Volcano plots displaying MAGeCK gene level enrichment score and log2 fold change for each gene tested in the secondary gRNA library, under lipoprotein-rich (A) or lipoprotein-depleted (D) cultured conditions, with genes identified with FDR<5% displayed in red. Venn diagrams demonstrating the overlap between HuH7 and HepG2 cells for positive (B, E) and negative (C, F) regulators of LDL uptake under lipoprotein-rich (B-C) or lipoprotein-depleted (E-F) culture conditions. (G) Positive regulators of LDL uptake in HuH7 cells under lipoprotein-depleted conditions were grouped by quartile and the proportion in each group that also influenced LDL uptake in HepG2 cells is displayed. (H) The effect size in lipoprotein-depleted conditions for gene-level gRNA enrichment in each cell type is plotted for genes showing a functional role in either cell type.

## DISCUSSION

Forward genetic screens are a powerful tool for the high-throughput and unbiased identification of genes that contribute to a biologic phenotype. Over the past decade, breakthroughs in genome editing technology have revolutionized the interrogation of gene function by improving the ease, speed, and accuracy of gene disruption. The programmability of CRISPR-mediated genome editing with a short gRNA sequence lends itself to large-scale oligonucleotide synthesis and quantification through massively parallel DNA sequencing. Together, these features make pooled CRISPR screening a powerful recent addition to the biologist’s toolkit.

We applied genome-wide CRISPR screening to identify novel determinants of cellular LDL uptake, identifying a large set of genes, many of which were not previously recognized to play a role in LDL uptake. The validity of our results is supported by several lines of evidence. First, we identified several well-established genes involved in cellular LDL uptake, with *LDLR* and *MYLIP* representing the top hits for positive and negative regulation of LDL uptake in both HuH7 and HepG2 cells, under both lipoprotein-rich and lipoprotein-depleted conditions, as well as for positive and negative regulation of cell surface and total LDLR abundance in HuH7 cells. Additional genes consistently identified across our screens included the positive control genes *SCAP*, *MBTPS1*, and *MBTPS2*. Second, our validation rate of hits was highly dependent on the strength of signal for a candidate gene in the primary screen, demonstrating a significant correlation over independent experiments. Third, our screen hits exhibited a high degree of concordance between lipoprotein-rich and lipoprotein-depleted conditions, much greater than might be expected by stochastic variation alone. This concordance also extended to the individual gRNA level, as gRNAs showing significant activity for one condition were much more likely to show activity for the other condition. Finally, we also observed a high degree of concordance between orthogonal screens. For example, genes whose perturbation impacted LDL uptake were much more likely to also be associated with reduced surface or total LDLR abundance.

Our findings highlight the value of following up candidate genes from a primary genome-wide CRISPR screen with a customized gRNA library. A more limited gene list allows for greater depth of gRNA per gene, infected cells per gRNA, and sequencing reads per gRNA. Reflecting these technical advantages, we observed significantly improved signal-to-noise ratio in our secondary screen, with more significant enrichment for positive hits, improved detection of internal positive controls, and no identification of negative control nontargeting genes. Generation of a secondary library also facilitates additional assays providing further biologic insight into screen hits, as we were more readily able to query candidate genes under different selective pressures.

Despite these strengths, a number of caveats apply to our screen data. First, our screen was performed in immortalized hepatoma cells, removed from the *in vivo* environment and evaluated in two-dimensional cell culture. While it is reassuring that this system recapitulates the LDLR-dependent, SREBP-responsive nature of cellular LDL uptake, the extent to which these interactions extend to the physiologic setting remains uncertain. The high degree of cell type specificity for our screen hits emphasizes the need for empirical testing of identified genes in other contexts. Second, the threshold for determining what constitutes a valid result is somewhat arbitrary. It is likely that among our list of hits are a subset of false positives, and likewise that among our genes which did not pass validation are a number of false negatives. Third, our screen may not uncover genes truly involved in LDL uptake if those genes also are either essential or confer a fitness advantage in culture, since gRNAs targeting those genes will be progressively depleted from the pooled population over the duration of the experiment. Fourth, our screen is unable to detect genes that perform redundant functions in LDL uptake, as compensation may prevent a significant phenotypic effect. For example, despite their clear roles in LDLR expression, we did not detect significant effects upon disruption of *SREBF1* or *SREBF2*, likely due to overlapping functions allowing one gene to compensate for loss of the other[7]. Finally, our screen is limited in detecting only those genes which exhibit a phenotype through a cell-autonomous effect. For example, PCSK9 induces LDLR degradation after its secretion. Therefore, *PCSK9*-targeted cells are still susceptible to the activity of PCSK9 secreted by neighboring cells, preventing these mutants from developing alterations in LDLR abundance and LDL uptake.

Orthogonal testing of our customized gRNA library provided us with initial insight into the mechanism of effect for each of our screen hits. Disruption of most LDL uptake regulators did not cause a similar reduction in transferrin uptake. Rather we frequently observed an inverse relationship, with some mutants exhibiting deficient uptake for one ligand and enhanced uptake for the other ligand. This pattern is consistent with distinct pathways of clathrin-mediated endocytosis and recycling of transferrin receptor and LDL receptor, as supported by findings from several groups[18–22]. Our findings suggest that perturbation of one pathway may indirectly cause an inverse effect on the other.

Screening of LDL uptake in a *LDLR*-deleted clone confirmed that many of our screen hits were dependent on *LDLR* for their functional influence on LDL uptake. Several LDL uptake modifiers however also seemed to influence LDL uptake on a *LDLR*-deleted genetic background. The molecular basis of this residual LDL uptake in LDLR-deleted cells is not well understood, but may be mediated by alternative receptors for LDL. For example, *SCARB1* encodes a scavenger receptor that binds a variety of ligands including LDL, has SREBP-binding sites in its promoter region, and is expressed in hepatocytes[23]. Supporting this model, we found disruption of *SCARB1* to be associated with reduced LDL uptake in both WT and *LDLR*-deleted HuH7 cells. LDL uptake regulators may therefore still contribute to LDL uptake on a *LDLR*-deleted background if they also influence *SCARB1* or other pathways mediating this residual LDL uptake.

We found that many LDL uptake regulators did not exhibit a readily detectable impact on LDLR levels either at the cell surface or associated with the entire cell, despite the apparent specificity of this antibody-based detection, with *LDLR* and *MYLIP* representing the top positive and negative regulators for each screen. The basis for this discrepancy is unclear but may be related either to screen hits influencing LDLR kinetics or function rather than steady state levels, or to compensatory effects in mutant cells that upregulate *LDLR* expression in response to defective LDL uptake.

Functional annotations of our novel screen hits showed modest enrichment in some pathways, including N-glycosylation, ubiquitination, and transcriptional regulation. In addition, the regions containing the identified genes were enriched for significant associations with LDL in a genome-wide association study of nearly 400,000 Europeans. Our findings provide further support for the involvement of these genes in human cholesterol regulation and suggest a molecular mechanism for their involvement in human lipid traits.

In summary, we identified a list of high-confidence genetic modifiers of HuH7 cell LDL uptake, with supporting evidence for their specificity, mechanism of action, and generalizability. These findings highlight the power of genome-scale CRISPR screening and offer new avenues for understanding the molecular determinants of cellular LDL uptake.

---

**Supplemental Figure 1.**
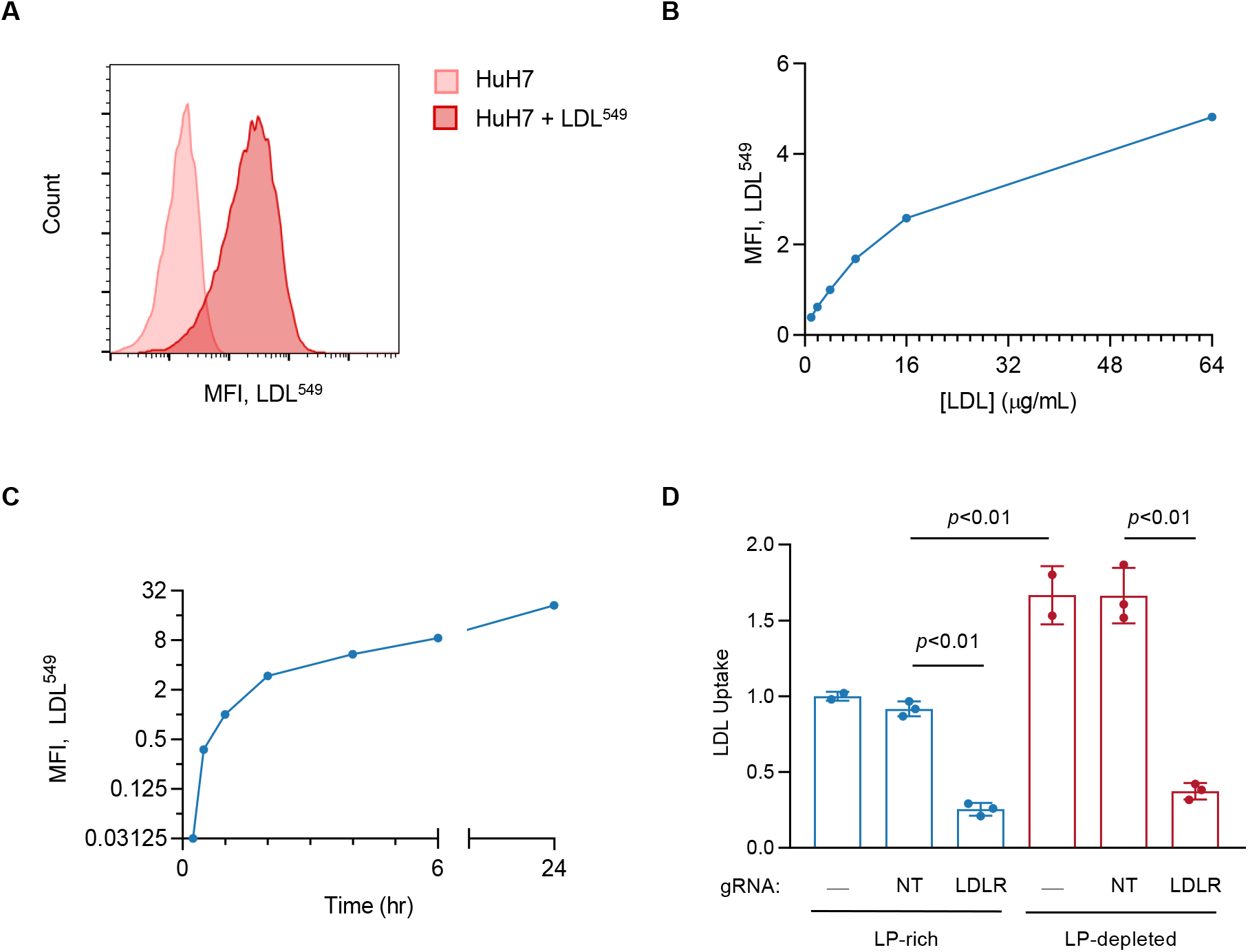
Development of conditions for primary screen of cellular LDL uptake. (A) Flow cytometry of HuH7 cells incubated for 1 hr in serum-free media with 4 μg/mL DyLight549-conjugated LDL, compared to autofluorescence of untreated HuH7 cells. (B) Dose-response curve of fluorescent signal acquisition by HuH7 cells incubated with a range of concentrations of DyLight549-conjugated LDL. (C) Time course of uptake of 4 μg/mL DyLight549-conjugated LDL by HuH7 cells. (D) Relative uptake was quantified by flow cytometry for WT HuH7 cells and cells targeted by CRISPR with a LDLR-targeting gRNA, or a nontargeting control gRNA, in cells that were pre-treated for 24 with lipoprotein-depleted media or maintained in lipoprotein-rich media.

**Supplemental Figure 2.**
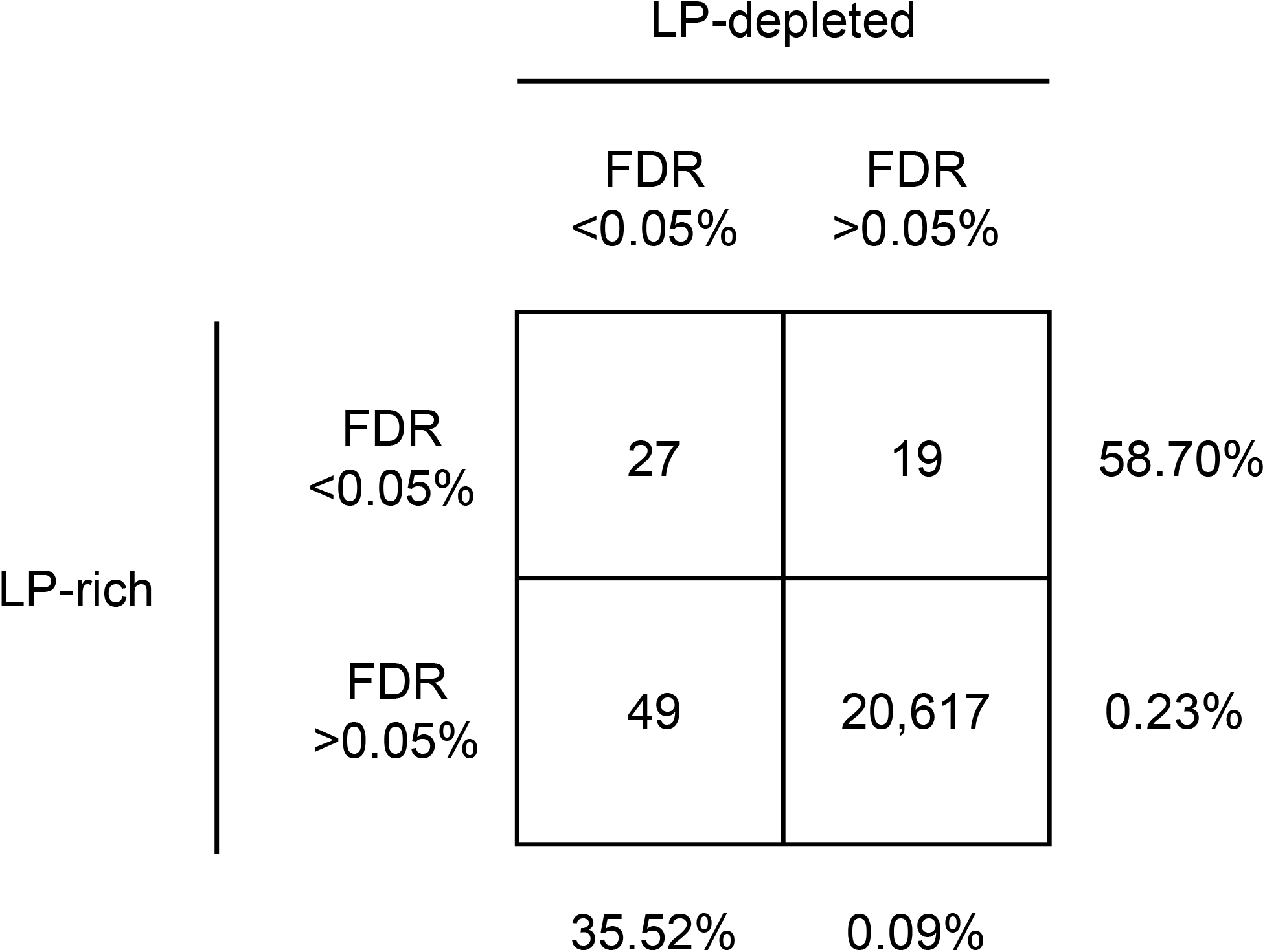
Concordance between HuH7 LDL uptake primary screen hits. The number of genes identified as positive regulators (FDR<0.05) under lipoprotein-rich and/or lipoprotein-depleted culture conditions is displayed.

**Supplemental Figure 3.**
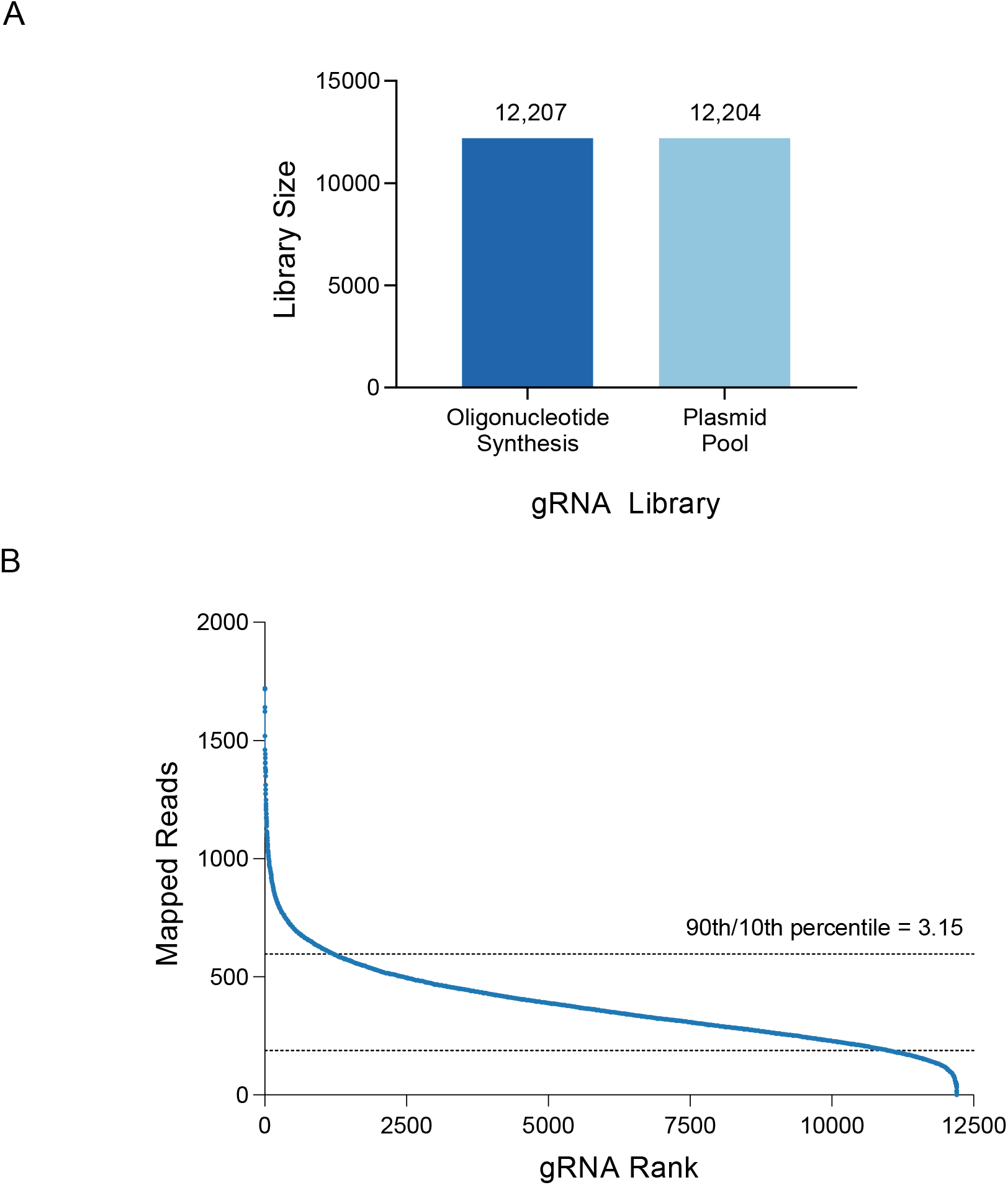
Synthesis of a customized gRNA library targeting candidate HuH7 LDL uptake regulators. (A) The number of unique gRNA sequences among the starting pooled oligonucleotide template and the synthesized plasmid pool are shown. (B) The number of mapped sequencing reads for each gRNA as a function of its relative rank in representation among all gRNAs. The ratio of reads for the gRNA at the 90^th^ and 10^th^ percentiles of representation are shown.

**Supplemental Figure 4.**
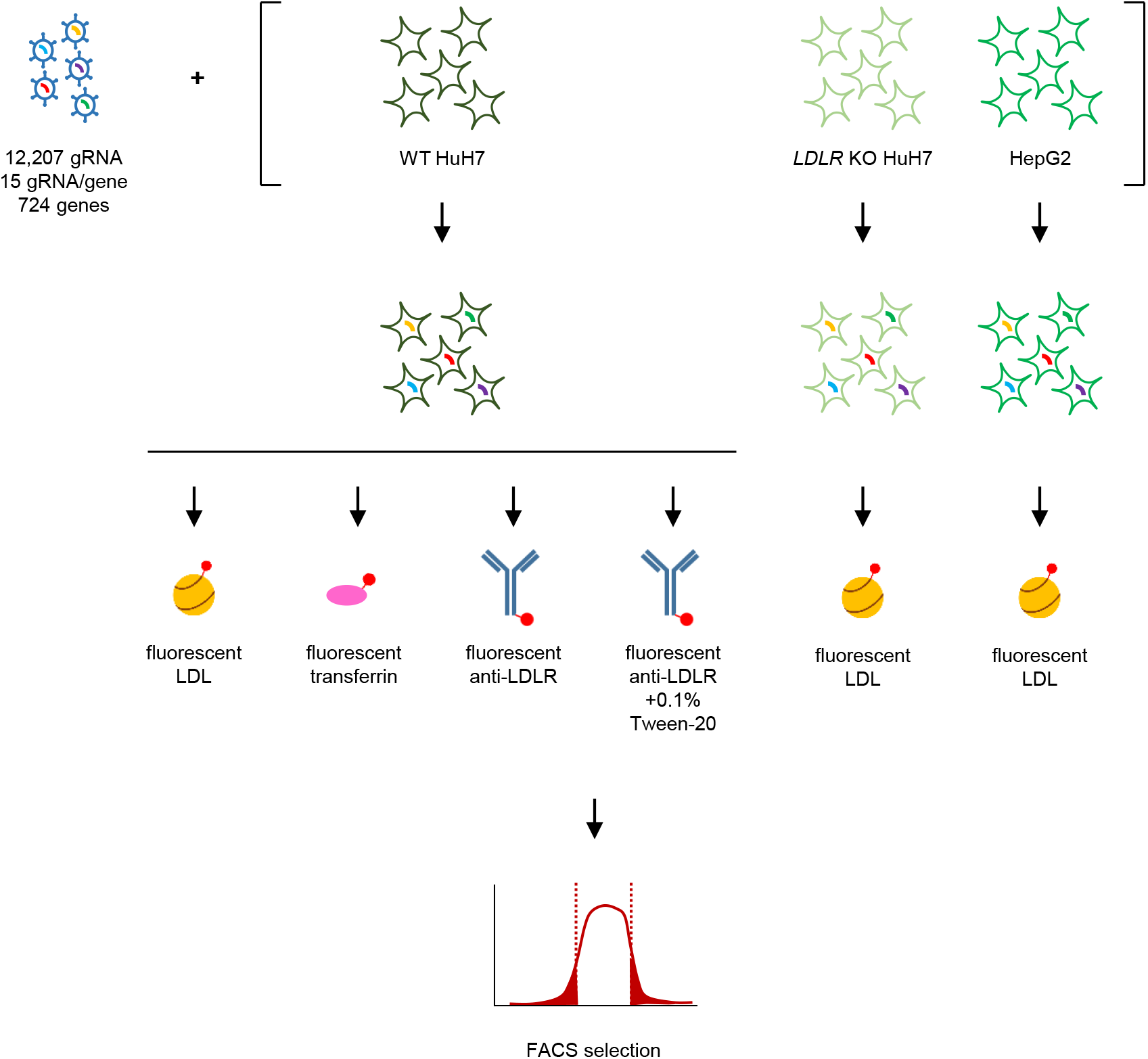
Strategy for secondary CRISPR validation screen and orthogonal screens. Mutagenesis of HuH7 WT cells, HuH7 LDLR KO cells, or HepG2 cells with the customized gRNA library is performed and pooled populations of mutants undergo selection by flow cytometry on the basis of relative LDL uptake, transferrin uptake, surface LDLR staining, or total cellular LDLR staining. The frequency of each gRNA in cells with high or low fluorescence is assessed by massively parallel DNA sequencing of gRNA amplicons, with computational analysis performed using the MAGeCK algorithm.

**Supplemental Figure 5.**
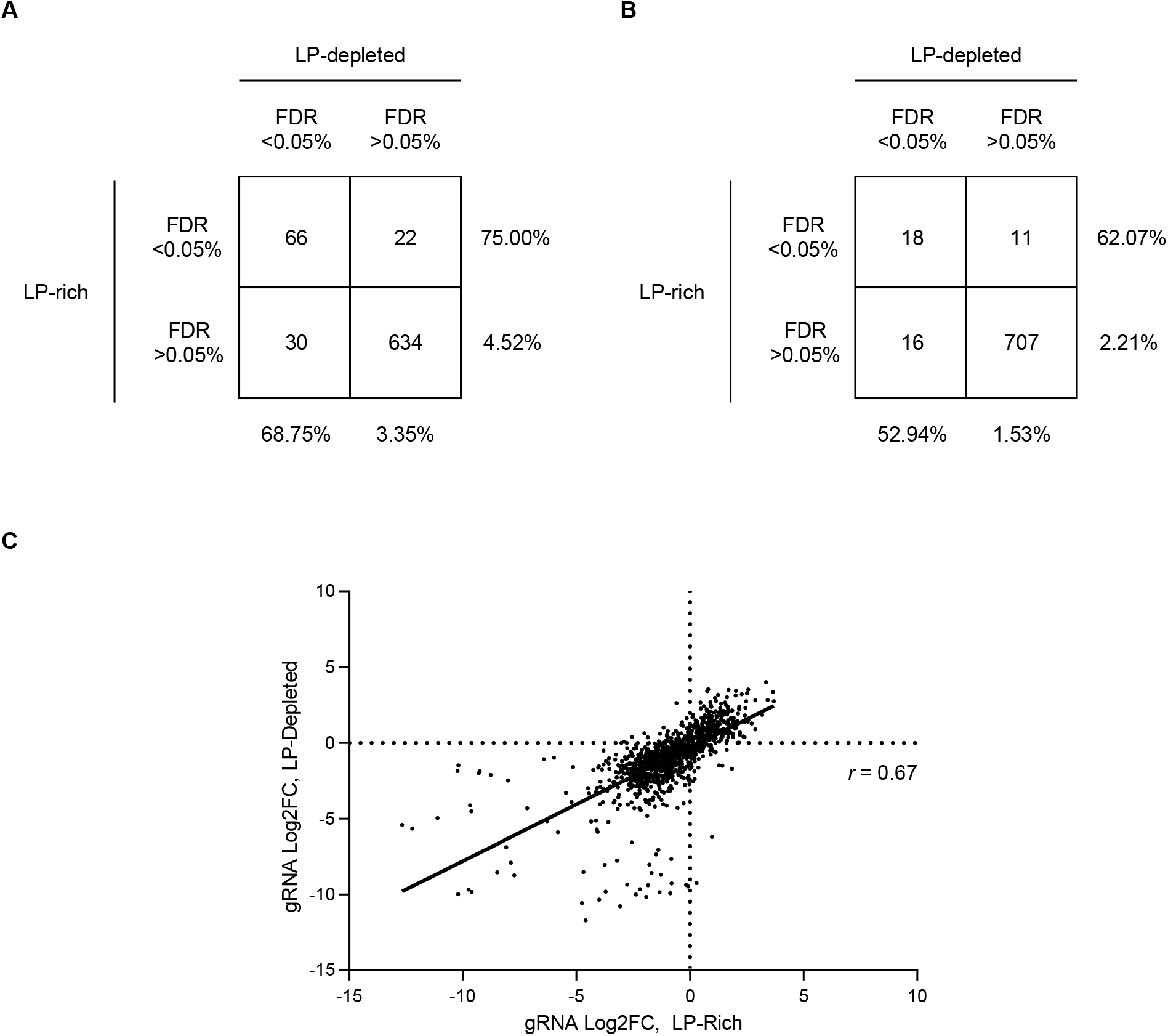
Concordance between HuH7 LDL uptake secondary screen hits. (A) For the secondary screen of HuH7 LDL uptake, the number of genes identified with a FDR<0.05 or FDR>0.05 under lipoprotein-rich and lipoprotein-depleted culture conditions is displayed. (B) Correlation between the degree of enrichment under lipoprotein-rich or lipoprotein-depleted conditions for each of the 15 gRNA for every target gene validated under both conditions by gene-level analysis.

**Supplemental Figure 6.**
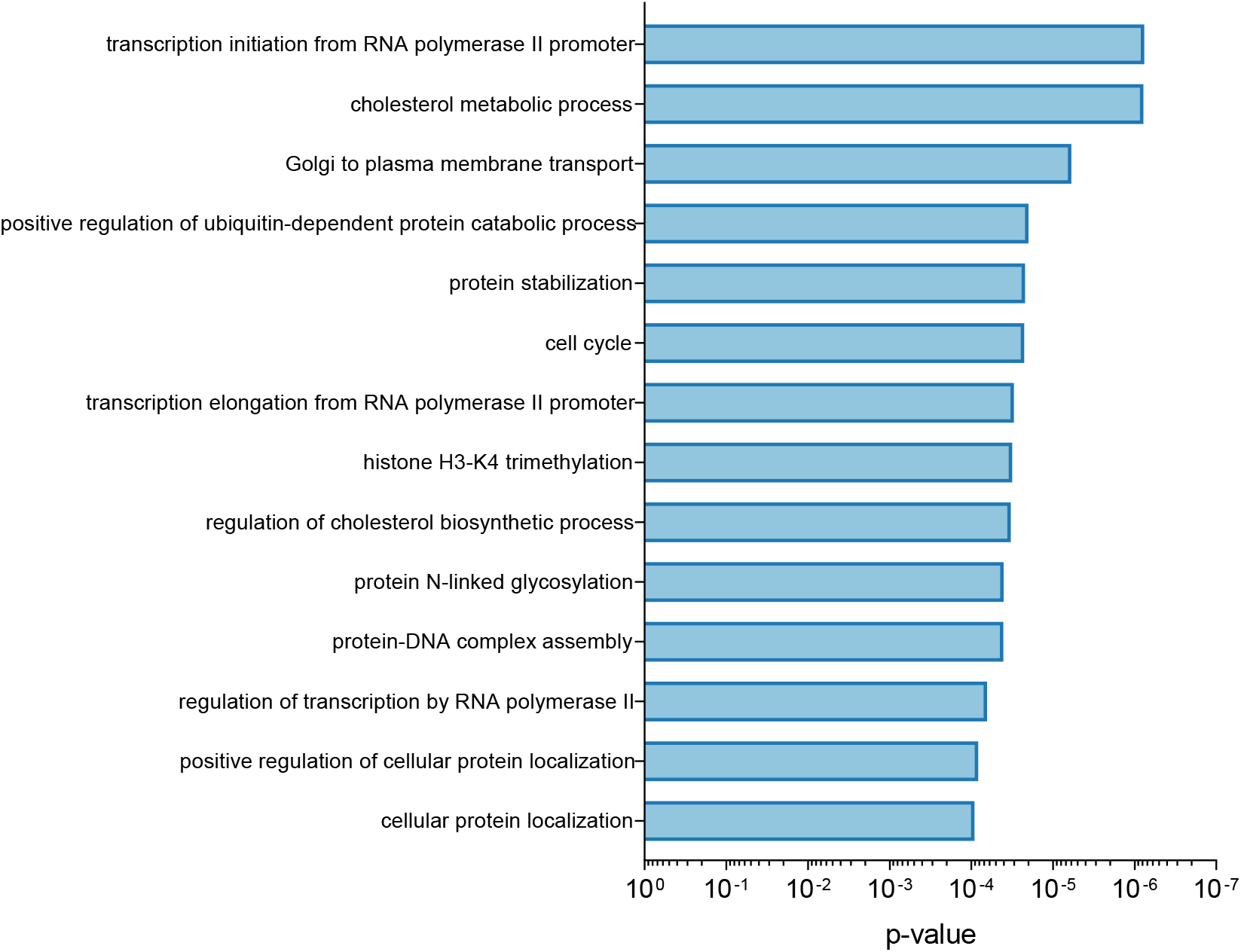
Functional annotations of validated LDL uptake regulators. Genes whose disruption was associated with a significant increase or decrease in LDL uptake, under either lipoprotein-rich or lipoprotein-depleted conditions, were analyzed by Gene Ontology classifications. Annotations demonstrating an enrichment with p<10^−4^ are displayed. Parental classifications for each are omitted from this figure and included in Supplemental Table 3.

**Supplemental Table 1.** MAGeCK analysis of primary genome-wide CRISPR screen of HuH7 LDL uptake.

**Supplemental Table 2.** MAGeCK analysis of targeted secondary CRISPR screens for modifiers of LDL uptake by HuH7 cells.

**Supplemental Table 3.** Functional annotation of HuH7 LDL uptake regulators identified in this study.

**Supplemental Table 4.** Significant UK Biobank LDL GWAS results within and nearby each gene

**Supplemental Table 5.** MAGeCK analysis of orthogonal CRISPR screen for modifiers of transferrin uptake by HuH7 cells.

**Supplemental Table 6.** MAGeCK analysis of orthogonal CRISPR screen for modifiers of LDL uptake by LDLR-deleted HuH7 cells.

**Supplemental Table 7.** MAGeCK analysis of orthogonal CRISPR screen for modifiers of LDLR abundance in HuH7 cells.

**Supplemental Table 8.** Comparison of genes identified in a previous siRNA screen of LDL uptake by endothelial cells and in this screen.

**Supplemental Table 9.** MAGeCK analysis of orthogonal CRISPR screen for modifiers of LDL uptake by HepG2 cells.

## MATERIALS AND METHODS

### Reagents

HuH7 cells were cultured in DMEM containing 10% FBS and penicillin/streptomycin. Cellular uptake assays were performed with fluorescent conjugates of LDL (Cayman Chemical, Ann Arbor MI, 10011229) or transferrin (ThermoFisher Scientific, Waltham MA, T35352). For immunoblotting, membranes were probed with antibodies against LDLR (Abcam, Cambridge UK, ab52818, 1:2000) and β-actin (Santa Cruz Biotechnology, Dallas TX, sc-47778, 1:5000). For flow cytometry, cells were stained with a fluorescently-conjugated antibody against LDLR (R&D Systems, Minneapolois MN, FAB2148G). CRISPR-mediating *LDLR* disruption was performed by cloning the gRNA sequence [AACAAGTTCAAGTGTCACAG] into BsmBI sites of pLentiCRISPRv2 (Addgene #52961, a gift from Feng Zhang[16]) or BbsI sites of pX459 (Addgene #62988, a gift from Feng Zhang). The nontargeting gRNA sequence [CGTGTGTGGGTAAACGGAAA] was used as a negative control. Genotyping primers for PCR amplification and Sanger sequencing of the LDLR targeting site were forward primer [TCCCAAAGTGCTGGGATTAC] and reverse primer [GGCAGAGTGGAGTTCCCAAA].

### Primary screen of HuH7 cellular LDL uptake

For each biologic replicate, 62.5 million HuH7 cells were harvested and evenly distributed into 12 separate 15 cm^2^ tissue culture plates. Pooled lentivirus containing the GeCKOv2 library[16] was added to cells in suspension at an estimated MOI of 0.4. The following day puromycin was added at a concentration of 1 μg/mL to select for infected cells. Cultured cells were then harvested, pooled, and passaged as needed to maintain logarithmic phase growth. Total cell numbers were maintained above 25 million cells (representing 200X coverage of the gRNA library) throughout the entirety of the screen. On assay day 12, cells were split into duplicate plates. On day 13, cells were either maintained in lipoprotein-rich media, or the media exchanged to DMEM supplemented with 10% lipoprotein-depleted fetal calf serum (Sigma S5394). On day 14, plates were sequentially processed by aspiration of media, washing in PBS, and addition of serum-free DMEM containing 4 μg/mL DyLight549-conjugated human LDL (Cayman Chemical, Ann Arbor MI, 10011229). Cells were incubated for 1 hr at 37°C then harvested with trypLE express, centrifuged 500×*g* for 5 min, washed in PBS, centrifuged again, resuspended at 20 million cells/mL PBS, and filtered into FACS tubes. Cell suspensions were then analyzed on a BD FACSAria III with cells exhibiting the top and bottom 75% DyLight^549^ fluorescence sorted into separate collection tubes. Genomic DNA was isolated using a DNEasy DNA isolation kit (Qiagen, Hilden, Germany). Preparation of barcoded amplicon libraries and mapping and deconvolution of sequencing reads obtained from an Illumina NextSeq sequencing run were performed as previously described[14]. Enrichment analysis was performed using the MAGeCK software package[24].

### Design and synthesis of secondary CRISPR library

Candidate genes from the primary LDL uptake screen were sorted by their relative ranking for MAGeCK gene level enrichment score. An FDR cutoff of 50% and 75% was used to select candidate positive and negative regulators, respectively. The candidate gene list was entered into the Broad Genetic Perturbation Platform sgRNA Designer for selection of 15 optimized targeting sequences per gene[25]. Nontargeting controls, long non-coding RNA, and microRNA candidates for which a corresponding target sequence could not be readily identified in the GPP platform were omitted. A total of 12 additional genes serving as internal controls (e.g. *TFRC* for transferrin uptake) or hypothesis-driven candidates (e.g. *SREBF2* for LDL uptake) were manually added to the candidate gene lists. Flanking sequences were appended to gRNA sequences to serve as priming sites for PCR amplification. Synthesized pooled oligonucleotides were obtained from CustomArray (Bothell, WA), amplified 18 cycles with Herculase II DNA polymerase (Agilent, Santa Clara CA), and purified using a QIAquick PCR purification kit (Qiagen). Assembly was performed with 1650 ng of BsmBI-digested pLentiCRISPRv2 and 250 ng of amplicon in a total reaction volume of 100 μL with HiFI DNA Assembly Mix (NEB, Ipswich MA) for 30 min at 50°C. Assembly products were purified with a QIAquick PCR purification kit, electroporated in triplicate into Endura electrocompetent cells (Lucigen, Middleton WI), and plated onto 24.5 cm^2^ LB-agar plates. After 14 hr at 37°C, bacteria were harvested and plasmid DNA purified with an EndoFree Plasmid Maxi kit (Qiagen). Dilution plates of electroporated cells confirmed a colony count of >100X relative to the size of the gRNA library. Library diversity was assessed with a Illumina MiSeq run of gRNA amplicons prepared as previously described[14].

### Validation and orthogonal screening of LDL uptake modifiers

Lentiviral infection with the customized CRISPR library, selection of infected cells, passaging, and parameters for LDL uptake were performed as in the primary genome-wide CRISPR screen. Transferrin uptake was performed with 5 μg/mL AlexaFluor^555^-conjugated transferrin (ThermoFisher) in serum-free DMEM for 30 min at 37°C. LDLR staining was performed for 30 min on ice with a 1:50 dilution of AlexaFluor^488^-conjugated LDLR antibody (R&D Systems, Minneapolois MN, FAB2148G) into PBS supplemented with 1% FBS, with or without 0.1% Tween-20 for surface or total cellular staining, respectively. Treated cells were sorted into high and low populations of fluorescence, genomic DNA isolated, and gRNA sequencing performed as in the primary screen. Three replicates were performed for each screen. Cell numbers were maintained above a minimum depth of coverage of 500X relative to the customized gRNA library throughout the screen until the time of sorting. For each sort, approximately 10-20 million cells were analyzed with 1-2 million cells collected per population.

### Generation of *LDLR*-deleted HuH7 clone

HuH7 cells were transfected with a *LDLR*-targeting CRISPR pX459 construct using Lipofectamine LTX (ThermoFisher). After puromycin selection of transfected cells was complete, serial dilutions of cells were plated into 96 well plates. Wells containing a single colony of growth were then selected for expansion. Single cell clones were analyzed by genotyping at the *LDLR* target site and immunoblotting with antibodies against LDLR and β-actin.

### Comparison to GWAS lipid trait associations

Association analysis for LDL cholesterol was performed using SAIGE[26] for 388,629 individuals in the white British subset of UK Biobank[27]. Inverse-normalized residuals for LDL after adjustment for batch, principle components 1-4, age, and age^2 were generated separately in males and females and then combined. Pre-treatment LDL levels were estimated for individuals on lipid-lowering medication by dividing the measured LDL value by 0.7. Control genes for comparison with the experimentally identified genes were selected based on nearest matching for both total gene length and total exon length. Gene transcription and exon start and end positions were taken from the refFlat file provided by the USCS genome annotation database[28]. Genes that overlapped within 500 kb of the identified gene start and end positions were excluded from the pool of control genes prior to matching. Significance for the difference in distribution of GWAS result p-values between the identified genes and selected control genes was determined using a two-sided Kolmogorov-Smirnov test.

### Functional annotation of LDL uptake modifiers

A total of 163 genes for which targeting in the secondary CRISPR screen was associated with either an increase or decrease in LDL uptake with FDR<5%, under either lipoprotein-rich or lipoprotein-depleted conditions, were included for analysis. This gene list was queried for enrichment of Gene Ontology classifications relative to all genes in the reference human genome using the PANTHER statistical overrepresentation test (PANTHER version 15.0, release February 14, 2020)[30]. Complete results are given in Supplemental Table 3. Classifications with p-value <10^−4^ at the most terminal node in the hierarchy for each subgroup are displayed in Supplemental Figure 6.

## Acknowledgements

This research has been conducted using the UK Biobank Resource under application number 24460.

## Notes

### Competing Interest Statement

The authors have declared no competing interest.

## REFERENCES

1. Ference BA, Ginsberg HN, Graham I, Ray KK, Packard CJ, Bruckert E, et al. Low-density lipoproteins cause atherosclerotic cardiovascular disease. 1. Evidence from genetic, epidemiologic, and clinical studies. A consensus statement from the European Atherosclerosis Society Consensus Panel. Eur Heart J. 2017;38(32):2459–72. Epub 2017/04/27. doi: 10.1093/eurheartj/ehx144. PubMed PMID: 28444290; PubMed Central PMCID: PMCPMC5837225.

2. Luo J, Yang H, Song BL. Mechanisms and regulation of cholesterol homeostasis. Nat Rev Mol Cell Biol. 2019. Epub 2019/12/19. doi: 10.1038/s41580-019-0190-7. PubMed PMID: 31848472.

3. Goldstein JL, Brown MS. A century of cholesterol and coronaries: from plaques to genes to statins. Cell. 2015;161(1):161–72. Epub 2015/03/31. doi: 10.1016/j.cell.2015.01.036. PubMed PMID: 25815993; PubMed Central PMCID: PMCPMC4525717.

4. Kathiresan S, Srivastava D. Genetics of human cardiovascular disease. Cell. 2012;148(6):1242–57. Epub 2012/03/20. doi: 10.1016/j.cell.2012.03.001. PubMed PMID: 22424232; PubMed Central PMCID: PMCPMC3319439.

5. Horton JD, Cohen JC, Hobbs HH. Molecular biology of PCSK9: its role in LDL metabolism. Trends Biochem Sci. 2007;32(2):71–7. Epub 2007/01/12. doi: 10.1016/j.tibs.2006.12.008. PubMed PMID: 17215125; PubMed Central PMCID: PMCPMC2711871.

6. Rader DJ, Cohen J, Hobbs HH. Monogenic hypercholesterolemia: new insights in pathogenesis and treatment. J Clin Invest. 2003;111(12):1795–803. Epub 2003/06/19. doi: 10.1172/JCI18925. PubMed PMID: 12813012; PubMed Central PMCID: PMCPMC161432.

7. Horton JD, Goldstein JL, Brown MS. SREBPs: activators of the complete program of cholesterol and fatty acid synthesis in the liver. J Clin Invest. 2002;109(9):1125–31. Epub 2002/05/08. doi: 10.1172/JCI15593. PubMed PMID: 11994399; PubMed Central PMCID: PMCPMC150968.

8. Brown MS, Goldstein JL. The SREBP pathway: regulation of cholesterol metabolism by proteolysis of a membrane-bound transcription factor. Cell. 1997;89(3):331–40. Epub 1997/05/02. PubMed PMID: 9150132.

9. Willer CJ, Schmidt EM, Sengupta S, Peloso GM, Gustafsson S, Kanoni S, et al. Discovery and refinement of loci associated with lipid levels. Nat Genet. 2013;45(11):1274–83. Epub 2013/10/08. doi: 10.1038/ng.2797. PubMed PMID: 24097068; PubMed Central PMCID: PMCPMC3838666.

10. Paththinige CS, Sirisena ND, Dissanayake V. Genetic determinants of inherited susceptibility to hypercholesterolemia – a comprehensive literature review. Lipids Health Dis. 2017;16(1):103. Epub 2017/06/05. doi: 10.1186/s12944-017-0488-4. PubMed PMID: 28577571; PubMed Central PMCID: PMCPMC5457620.

11. Peloso GM, Natarajan P. Insights from population-based analyses of plasma lipids across the allele frequency spectrum. Curr Opin Genet Dev. 2018;50:1–6. Epub 2018/02/16. doi: 10.1016/j.gde.2018.01.003. PubMed PMID: 29448166; PubMed Central PMCID: PMCPMC6087690.

12. Dron JS, Hegele RA. Polygenic influences on dyslipidemias. Curr Opin Lipidol. 2018;29(2):133–43. Epub 2018/01/05. doi: 10.1097/MOL.0000000000000482. PubMed PMID: 29300201.

13. Shalem O, Sanjana NE, Zhang F. High-throughput functional genomics using CRISPR-Cas9. Nat Rev Genet. 2015;16(5):299–311. doi: 10.1038/nrg3899. PubMed PMID: 25854182; PubMed Central PMCID: PMCPMC4503232.

14. Emmer BT, Hesketh GG, Kotnik E, Tang VT, Lascuna PJ, Xiang J, et al. The cargo receptor SURF4 promotes the efficient cellular secretion of PCSK9. Elife. 2018;7. Epub 2018/09/27. doi: 10.7554/eLife.38839. PubMed PMID: 30251625; PubMed Central PMCID: PMCPMC6156083.

15. Nakabayashi H, Taketa K, Miyano K, Yamane T, Sato J. Growth of human hepatoma cells lines with differentiated functions in chemically defined medium. Cancer Res. 1982;42(9):3858–63. Epub 1982/09/01. PubMed PMID: 6286115.

16. Sanjana NE, Shalem O, Zhang F. Improved vectors and genome-wide libraries for CRISPR screening. Nat Methods. 2014;11(8):783–4. doi: 10.1038/nmeth.3047. PubMed PMID: 25075903; PubMed Central PMCID: PMCPMC4486245.

17. Zelcer N, Hong C, Boyadjian R, Tontonoz P. LXR regulates cholesterol uptake through Idol-dependent ubiquitination of the LDL receptor. Science. 2009;325(5936):100–4. Epub 2009/06/13. doi: 10.1126/science.1168974. PubMed PMID: 19520913; PubMed Central PMCID: PMCPMC2777523.

18. Lakadamyali M, Rust MJ, Zhuang X. Ligands for clathrin-mediated endocytosis are differentially sorted into distinct populations of early endosomes. Cell. 2006;124(5):997–1009. Epub 2006/03/15. doi: 10.1016/j.cell.2005.12.038. PubMed PMID: 16530046; PubMed Central PMCID: PMCPMC2660893.

19. Bartuzi P, Billadeau DD, Favier R, Rong S, Dekker D, Fedoseienko A, et al. CCC- and WASH-mediated endosomal sorting of LDLR is required for normal clearance of circulating LDL. Nat Commun. 2016;7:10961. Epub 2016/03/12. doi: 10.1038/ncomms10961. PubMed PMID: 26965651; PubMed Central PMCID: PMCPMC4792963.

20. Keyel PA, Mishra SK, Roth R, Heuser JE, Watkins SC, Traub LM. A single common portal for clathrin-mediated endocytosis of distinct cargo governed by cargo-selective adaptors. Mol Biol Cell. 2006;17(10):4300–17. Epub 2006/07/28. doi: 10.1091/mbc.e06-05-0421. PubMed PMID: 16870701; PubMed Central PMCID: PMCPMC1635374.

21. Brown MS, Anderson RG, Goldstein JL. Recycling receptors: the round-trip itinerary of migrant membrane proteins. Cell. 1983;32(3):663–7. Epub 1983/03/01. doi: 10.1016/0092-8674(83)90052-1. PubMed PMID: 6299572.

22. Mayle KM, Le AM, Kamei DT. The intracellular trafficking pathway of transferrin. Biochim Biophys Acta. 2012;1820(3):264–81. Epub 2011/10/05. doi: 10.1016/j.bbagen.2011.09.009. PubMed PMID: 21968002; PubMed Central PMCID: PMCPMC3288267.

23. Shen WJ, Asthana S, Kraemer FB, Azhar S. Scavenger receptor B type 1: expression, molecular regulation, and cholesterol transport function. J Lipid Res. 2018;59(7):1114–31. Epub 2018/05/04. doi: 10.1194/jlr.R083121. PubMed PMID: 29720388; PubMed Central PMCID: PMCPMC6027903.

24. Li W, Xu H, Xiao T, Cong L, Love MI, Zhang F, et al. MAGeCK enables robust identification of essential genes from genome-scale CRISPR/Cas9 knockout screens. Genome Biol. 2014;15(12):554. doi: 10.1186/s13059-014-0554-4. PubMed PMID: 25476604; PubMed Central PMCID: PMCPMC4290824.

25. Sanson KR, Hanna RE, Hegde M, Donovan KF, Strand C, Sullender ME, et al. Optimized libraries for CRISPR-Cas9 genetic screens with multiple modalities. Nat Commun. 2018;9(1):5416. Epub 2018/12/24. doi: 10.1038/s41467-018-07901-8. PubMed PMID: 30575746; PubMed Central PMCID: PMCPMC6303322.

26. Zhou W, Nielsen JB, Fritsche LG, Dey R, Gabrielsen ME, Wolford BN, et al. Efficiently controlling for case-control imbalance and sample relatedness in large-scale genetic association studies. Nat Genet. 2018;50(9):1335–41. Epub 2018/08/15. doi: 10.1038/s41588-018-0184-y. PubMed PMID: 30104761; PubMed Central PMCID: PMCPMC6119127.

27. Sudlow C, Gallacher J, Allen N, Beral V, Burton P, Danesh J, et al. UK biobank: an open access resource for identifying the causes of a wide range of complex diseases of middle and old age. PLoS Med. 2015;12(3):e1001779. Epub 2015/04/01. doi: 10.1371/journal.pmed.1001779. PubMed PMID: 25826379; PubMed Central PMCID: PMCPMC4380465.

28. Karolchik D, Hinrichs AS, Kent WJ. The UCSC Genome Browser. Curr Protoc Bioinformatics. 2009;Chapter 1:Unit1 4. Epub 2009/12/04. doi: 10.1002/0471250953.bi0104s28. PubMed PMID: 19957273; PubMed Central PMCID: PMCPMC2834533.

29. Eden E, Navon R, Steinfeld I, Lipson D, Yakhini Z. GOrilla: a tool for discovery and visualization of enriched GO terms in ranked gene lists. BMC Bioinformatics. 2009;10:48. Epub 2009/02/05. doi: 10.1186/1471-2105-10-48. PubMed PMID: 19192299; PubMed Central PMCID: PMCPMC2644678.

30. Mi H, Muruganujan A, Ebert D, Huang X, Thomas PD. PANTHER version 14: more genomes, a new PANTHER GO-slim and improvements in enrichment analysis tools. Nucleic Acids Res. 2019;47(D1):D419–D26. Epub 2018/11/09. doi: 10.1093/nar/gky1038. PubMed PMID: 30407594; PubMed Central PMCID: PMCPMC6323939.

